# Single-nucleus RNA sequencing reveals cell type-specific responses to heat stress in bovine mammary gland

**DOI:** 10.64898/2026.02.18.706602

**Authors:** Xingtan Yu, Shambhvi, Diego A. Ceballos, Marina M. Ferreira, Alejandra Zapata, Nirosh Seneviratne, Sabina Pokharel, Yifei Fang, Guangsheng Li, Francisco Leal-Yepes, Joseph W. McFadden, Jingyue Ellie Duan

**Affiliations:** Department of Animal Science, College of Agriculture and Life Sciences, Cornell University, Ithaca, 14853, USA; Department of Population Medicine and Diagnostic Sciences, College of Veterinary Medicine, Cornell University, Ithaca, 14853, USA

**Keywords:** Dairy cattle, Mammary gland, Heat stress, Single-nucleus RNA sequencing

## Abstract

**Background:** Heat stress (HS) poses a major challenge to the dairy industry by reducing milk production, yet its cell type-specific effects in the bovine mammary gland remain incompletely defined. In this study, we recorded production traits and collected mammary biopsies from cows under thermoneutral (TN), HS, and pair-fed (PF) conditions.

**Results:** Clinical measurements confirmed HS-induced physiological alterations. Compared with TN cows, HS cows exhibited reduced dry matter intake (DMI), milk yield, and yields of fat, protein, and lactose, along with increased water intake and milk urea nitrogen. The use of PF controls indicated that decreased DMI accounted for 45% of the milk-yield reduction, whereas direct HS effects accounted for the remaining reduction.

We applied single-nucleus RNA-seq (snRNA-seq) on mammary biopsies to generate cell-resolved HS responses. We identified 14 distinct cell clusters, including epithelial, immune, and stromal populations. Under the TN condition, casein genes (e.g., *CSN1S1*, *CSN2*) were broadly expressed across luminal cells but were attenuated under HS, whereas luminal alveolar cells showed relative upregulation. Heat shock protein genes were strongly induced by HS, primarily in epithelial clusters. Gene-set enrichment analyses revealed increased ribosomal activities across HS-responsive clusters and enrichment of protein folding and metabolic pathways in luminal alveolar cells, suggesting elevated proteostasis demands under stress.

Pseudotime analysis positioned luminal cells along a progenitor-to-secretory trajectory under TN, accompanied by increased casein gene expression, whereas under HS, mature luminal cells shifted toward a homeostasis regulatory state. Cell-cell communication analysis demonstrated HS-induced remodeling of interepithelial signaling, including altered ERBB4-mediated signaling from luminal hormone-sensing to alveolar lineages. Finally, transcription factor activity profiling highlighted cell type-specific HS-activated regulators and their downstream target genes.

**Conclusions:** Together, this cell type-resolved atlas delineates how HS alters bovine mammary epithelial function, developmental state, and intercellular crosstalk. These findings point to proteostasis pressure, disrupted signaling pathways, and rewired regulatory networks as mechanistic contributors to reduced lactational performance under HS, offering insights for improving heat resilience in dairy cattle.

## Background

As global temperatures rise, heat stress has become a severe challenge for the dairy industry, compromising animal welfare and reducing productivity [1,2]. Heat stress disrupts cows’ thermal homeostasis and induces systemic physiological changes, including elevated body temperature, increased respiration rate, altered peripheral blood flow, and reduced dry matter intake (DMI) [3,4]. These changes generally impact lactation, causing notable decreases in milk production and composition, which ultimately lower overall feed efficiency [5]. Although reduced DMI contributes to these changes, pair-feeding studies showed that intake repression explains only 30-50% of the yield loss under heat stress [6,7], suggesting that HS directly impairs mammary gland (MG) function. Understanding these direct effects is essential for clarifying how thermal stress alters mammary cellular activity and constrains milk synthesis beyond systemic metabolic adjustments.

Bulk transcriptomic analyses of mammary tissue from heat-stressed cows have revealed downregulation of milk protein genes, including caseins, accompanied by upregulation of heat shock protein genes (HSPs). Enrichment of immune response and metabolism pathways has also been reported [8,9], indicating broad cellular transcriptomic changes under HS. However, the MG is a complex organ composed of heterogeneous cell populations. Mammary epithelial cells include both myoepithelial cells and luminal cells. Myoepithelial cells form the outer layer and contract to facilitate milk ejection from the inner luminal cells [10]. Luminal cells can be further divided into two lineages, hormone-sensing and alveolar, which respond to hormonal signals and synthesize milk, respectively [11]. Previous studies have also identified a rare population of luminal cells expressing markers of both lineages, indicating an undifferentiated luminal cell state [12]. In addition, stromal cells such as fibroblasts, endothelial cells, and immune cells all contribute to MG function and interact with epithelial cells, and each cell type is likely to respond differently to thermal stress [13]. Conventional bulk RNA sequencing masks cell type-specific transcriptional dynamics, leaving a gap in understanding how HS affects distinct MG cell populations.

Single-nucleus RNA sequencing (snRNA-seq) enables the high-resolution profiling of individual nuclei from complex tissues, revealing cellular diversity, developmental trajectories, intercellular communication, and regulatory networks [14]. Unlike single-cell RNA sequencing (scRNA-seq), which requires isolation of viable whole cells from fresh tissue, snRNA-seq can be performed on frozen or fixed tissues [15]. By capturing nuclear transcripts, snRNA-seq preserves the transcriptional landscape of all major cell types while minimizing artifacts from enzymatic dissociation or cell stress responses [16]. This technique has also been increasingly adopted in MG biology to study developmental dynamics, cellular functions, and ancestry-dependent variability in mice and humans [17–19]. However, to our knowledge, snRNA-seq has not yet been applied to the bovine MG.

In this study, we applied snRNA-seq to mammary biopsies from late-lactating cows housed in respiration chambers (RC) under thermoneutral (TN) and heat stress (HS) conditions. A pair-fed (PF) control group was included to match the DMI of HS cows to decouple nutritional effects from physiological changes. We identified diverse epithelial, stromal, and immune cells and revealed cell type-specific response to HS characterized by elevated proteostasis demand. HS also altered intercellular communication, shifted luminal cell states from secretory to stress-adaptive, and activated stress-related regulatory networks, collectively providing mechanistic insights into milk reduction. Overall, this work presents the first single-nucleus transcriptomic atlas of the bovine mammary gland under environmental stress, offering a high-resolution framework to understand mammary cellular plasticity and to inform strategies for improving heat resilience in dairy cattle.

## Methods

### Animals and sample collection

All experimental procedures were approved by the Cornell University Institutional Animal Care and Use Committee (#2022-0186). Nine multiparous pregnant lactating Holstein dairy cows (mean ± s.d.; 304 ± 30.94 days in milk, 186 ± 14.43 days carrying calf, and 27.2 ± 3.44 kg of milk) were enrolled in a randomized complete block design. Cows were allocated into three blocks of three cows each, balanced for milk yield, days in milk, and days pregnant. Cows were transported from the Cornell University Research Center (Dryden, NY) to Cornell University Large Animal Research and Teaching Unit (Ithaca, NY) and acclimated in tie-stall housing under thermoneutral conditions (22 ± 0.16 °C; 45 ± 0.05% relative humidity; THI = 68) for three days.

On the fourth day, cows within a block were randomly allocated to one of three climate-controlled respiration chambers (No Pollution Ltd., Leicester, UK) and underwent a 4-h chamber acclimation during which their eating, drinking, and lying behaviors were monitored. Following acclimation, cows were returned to the tie-stalls overnight and moved back to the chambers at 0700 h the following morning. The experiment began with a 24-h baseline measurement under thermoneutral conditions (THI = 68) in the chambers, after which cows entered a 3-d treatment period and were randomly assigned to one of three environmental treatments (**Fig. 1**): thermoneutrality (TN; n = 3), heat stressed (HS; n = 3), and thermoneutrality but pair-fed to match the feed intake of heat-stressed cows (PF; n = 3). Thermoneutral chambers were programmed at constant 22 °C and 45% relative humidity (THI = 68). The heat-stress chamber was programmed to follow a diurnal cycle, with temperature increased from 27 to 37 °C at 0700 h (RH 50%) and decreased from 37 to 27 °C at 1900 h (RH 45%), targeting a daytime THI of 82 (not exceeding 86) and a nighttime THI of 74 (**Additional file 1: Fig. 1**). The temperature-humidity index (THI) was calculated as follows [20]: THI = (1.8 × Temperature + 32) − [(0.55 − 0.0055 × Relative Humidity) × (1.8 × Temperature − 26)]. One cow in the PF group was ill and therefore removed from the experiment, leaving n = 2 in the PF treatment. Cows were fed twice daily at 0630 and 1930 h and milked twice daily at 0600 and 1900 h. Cows had ad libitum access to water and total mixed ration (TMR; DM basis: 44.20% corn silage, 8.87% haylage, and 44.20% concentrate) formulated to meet or exceed the requirements. During the treatment period, PF cows followed a predefined feeding protocol in which daily feed allowances were adjusted based on the percentage reduction in intake observed in the corresponding HS cow. The daily intake reductions for HS cows were calculated relative to their average intake during the three days preceding treatment, which served as the baseline, and applied to PF cows using their own baseline intake.

**Fig. 1.**
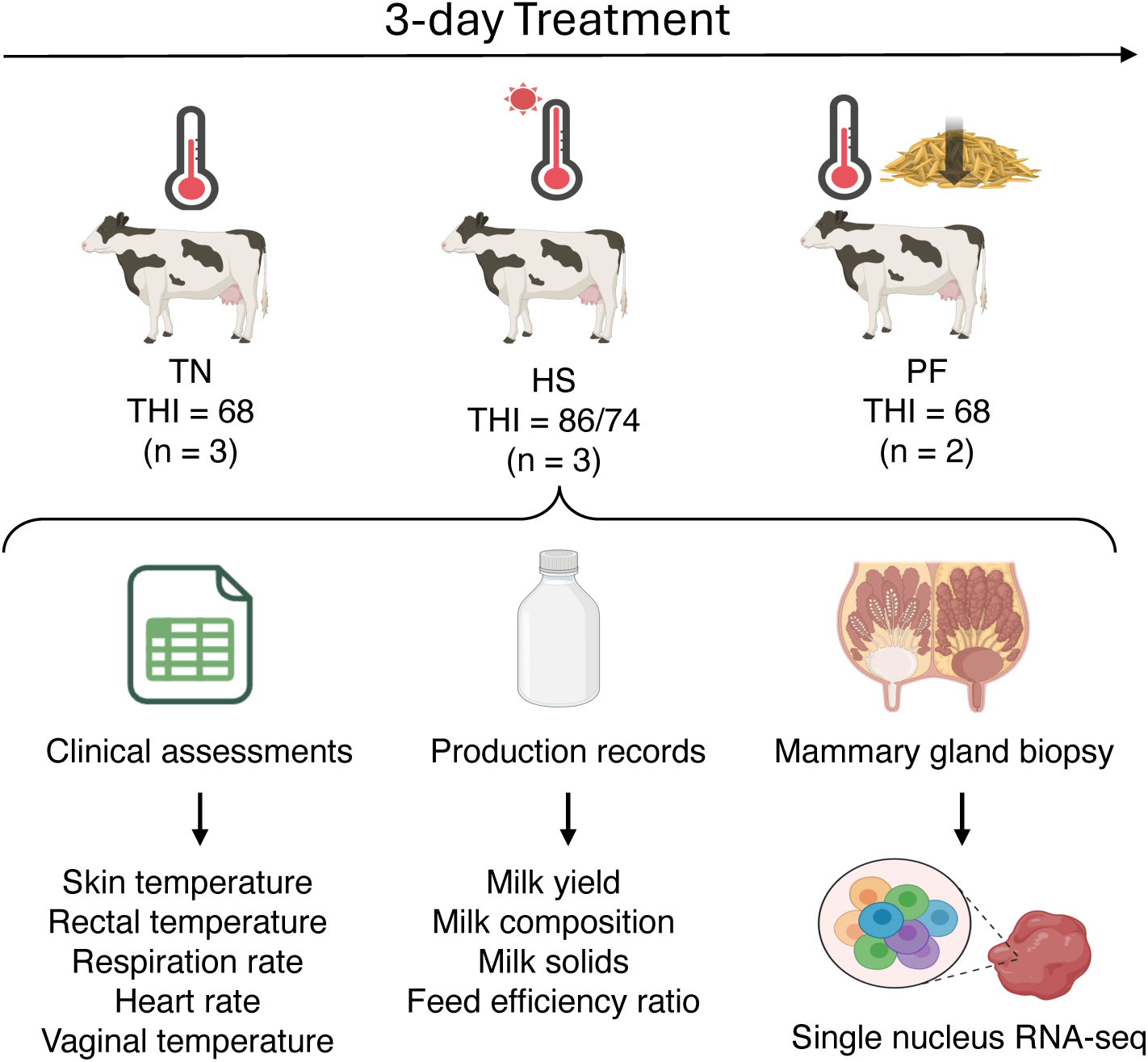
Experimental design and clinical assessments. Eight multiparous lactating Holstein cows were enrolled in the study for a 3-day treatment and randomly assigned into three conditions: thermoneutrality (TN, n = 3), heat stress (HS, n = 3), or thermoneutrality and pair-fed to HS (PF, n = 2). Clinical assessments and production records were collected during acclimation for baseline values and during treatment. Mammary biopsies were collected for single-nucleus RNA sequencing.

The next morning after the 3-d treatments, a mammary gland was collected. All cows were subject to a physical examination prior to the biopsy procedure. The mammary gland biopsy was performed following a validated methodology [21]. Briefly, all cows were restrained in a standing position, a dose of acepromazine (0.005 mg/kg) was administered via coccygeal vein, and flunixin meglumine was administered at a dose of 1.1 mg/kg via the jugular vein. All biopsies were collected from the right hind quarter (RH). The hair on the udder surface was clipped, and the biopsy area in the mammary gland was identified using ultrasound (IBEX PRO, E.I. Medical Imaging, Loveland, CO), avoiding large blood vessels and cisternal space. The area was scrubbed three times, alternating 4×4 gauze with povidone–iodine and 70% ethanol, followed by a bleb of 2% lidocaine and a final scrub repeating the same process previously described.

A 0.5-1 cm vertical skin incision was made with a #22 sterile scalpel blade, and a 2 mm disposable biopsy punch was inserted 0.5-1 cm into the mammary gland by applying pressure and rotating it clockwise. A catheter adaptor and 20 mL syringe were used as a suction device to ensure the tissue was extracted. The tissue was removed from the punch with sterile forceps, rinsed with 0.9% saline solution, and frozen in dry ice, then transported to the lab and stored at −80°C until nuclei preparation. The incision site was closed using a surgical skin stapler and AluSpray Aerosol Bandage was applied. Animals and incision sites were monitored daily until staple removal 7 d post-procedure.

### Physiological data collection and Modeling

Feed intake, water intake, and milk yield were recorded daily for the duration of the study. TMR and orts were collected every day during baseline and treatment period and analyzed for dry matter (AOAC International, 2000). Milk was sampled from 4 consecutive milkings prior to heat-stress exposure and from each milking throughout the treatment period. Samples were analyzed for fat, protein, lactose, and milk urea nitrogen (MUN) by Fourier-transform infrared spectroscopy (Milkoscan FT+; Foss Inc.) at Dairy One DHIA Laboratory (Ithaca, NY). Somatic cell count (SCC) was measured by flow cytometry (Fossomatic FC; Foss Inc.), and somatic cell score was calculated using the following logarithmic transformation: Log_2_ (somatic cell count/100,000) + 3 [22]. The yields of 3.5% fat corrected milk (FCM; [23], energy corrected milk (ECM; [24], and milk components were calculated based on milk yield and component concentrations and then summed to obtain daily totals. The efficiencies for milk yield, FCM (FCME), and ECM (ECME) production were calculated as the ratio of milk yield, FCM, or ECM in relation to dry matter intake.

Clinical parameters, including rectal temperature, skin temperature, heart rate, and respiration rate, were recorded three times daily at 0700, 1200, and 1900 h (**Additional file 2: Table S1**). Baseline values for parameters were calculated as the average of measurements obtained during the three days preceding the treatment period. Rectal temperature was measured using a digital large-animal thermometer (SHARPTEMP-V; Cortan Corp.). Skin temperature was obtained using a handheld infrared thermometer (model 568; Fluke Corp.) on a shaved area of the left flank. Respiration rate was determined by counting flank movements for 15 s and multiplying by four to calculate breaths per minute. Heart rate was measured using a stethoscope (model 10-404; MABIS Healthcare) placed caudal to the left elbow, with beats counted for 30 s and multiplied by two to obtain beats per minute.

Production and clinical data were analyzed using the following mixed model procedure in SAS (v9.4, SAS Institute Inc.):

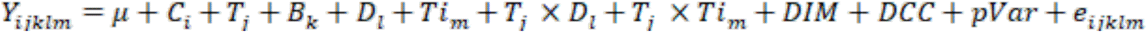

where Y_ijklm_ = dependent variable; μ = overall mean; C_i_ = random effect of cow (i = 1 to 8); T_j_ = fixed effect of treatment (j = 1 to 3); B_k_ = fixed effect of block (k = 1 to 3); D_l_ = fixed effect of day (l = 1 to 3); Ti_m_ = fixed effect of time (m = 1 to 3 for clinical parameters); T_j_ × D_l_ = fixed effect of the interaction between treatment and day; T_j_ × Ti_m_ = the fixed effect of the interaction between treatment and time; DIM = days in milk used as a covariate; DCC = days carrying calf used as a covariate; pVar = baseline measurement for each response variable used as a covariate; and e_ijklm_ = residual error. Model fit was assessed using variance components, first-order autoregressive, and compound symmetry covariance structures, and the structure with the smallest Bayesian Information Criterion (BIC) was selected. Model assumptions were assessed by inspecting residual distribution using probability and box plots and by examining variance patterns across predicted values. Preplanned non-orthogonal contrasts were used to compare the least squares means. Main effects and interactions were considered statistically significant at P ≤ 0.05 and P ≤ 0.15, respectively, with trends identified when 0.05 < P ≤ 0.15. Observations with studentized residuals > 3.0 or < −3.0 were considered outliers.

### Vaginal temperature measurement using HOBO devices and Analysis

Vaginal temperature was continuously monitored during the 3-d treatment using individual HOBO data loggers (Bourne, MA) programmed to record temperature at 1-min intervals. Progesterone intravaginal insert devices (CIDR; Zoetis, Parsippany, NJ) were preconditioned by soaking in water for 2-wk, with weekly water replacement, to minimize residual progesterone. The sensor tip of each HOBO data logger was affixed to the CIDR using vinyl electrical tape (Temflex; 3M, Austin, TX) and reinforced with elastic tape (Elastikon; Pinetown, South Africa). Immediately before insertion, the assembled device was disinfected in a 2% chlorhexidine solution. The vulva was cleaned with disposable paper towels, and the device was inserted using an intravaginal applicator. The HOBO logger was secured to the hip using veterinary livestock adhesive (TAG Cement; Ruscoe), allowing normal animal movement. The device was then activated using the HOBO Connect mobile application, and data were downloaded every 24 h. Devices were removed at the end of each cohort, and cows were monitored throughout the study for signs of discomfort or adverse health effects. Vaginal temperature was recorded every minute during the study. All temperatures below the physiological threshold (<37.9°C) were removed, and the average was calculated every 15 minutes for statistical analysis. Baseline was analyzed separately from the treatment period using PROC Mixed in SAS 9.4 and the Repeated statement. Fixed effects included in the model were treatment, time, and treatment*time interaction. Tukey’s post hoc test was used for multiple comparison correction of *P*-values for all pairwise comparisons of least squares means. Normality and homoscedasticity of residuals were tested for each model fit.

### Single nucleus extraction and sequencing

Frozen biopsies from the same treatment group were pooled into one sample and shipped to Novogene (Beijing, China) for single-nucleus isolation. Library preparation was performed using the 10x Genomics Chromium Single Cell 3’ v4 assay kit according to the manufacturer’s instructions. Briefly, cDNA was synthesized and amplified from barcoded nuclei and was then used to construct sequencing libraries. Library quality was assessed using an Agilent Bioanalyzer. Libraries were sequenced on an Illumina NovaSeq X Plus platform with paired-end 150 bp reads.

### Bioinformatics data analysis

10x snRNA sequence data were aligned to the bovine reference genome (ARS-UCD 1.2, Ensembl) using CellRanger (v9.0.1) [25] with default parameters. The output matrix was imported into R (v4.4.3), and cells expressing fewer than 200 genes and genes detected in fewer than three cells were filtered out. Subsequent quality control and visualization were performed using Seurat (v5.3.0) [26]. Cells with nCount_RNA between 1,000 and 40,000, nFeature_RNA between 200 and 2,500, and mitochondrial content below 10% were retained for analysis. This quality control step resulted in 5,201, 7,711, and 1,679 cells in TN, HS, and PF samples, respectively (**Additional file 3: Table S2**). The three datasets were integrated using the IntegrateData function with 5,000 variable genes. The optimal resolution for cell clustering was determined using the clustree (v0.5.1) package [27], and dimensionality reduction was performed using the top 15 principal components based on the elbow plot. The FindAllMarkers function was used to identify differentially expressed genes in each cell cluster. Genes were considered differentially expressed if the adjusted P value was < 0.05 and the log2(fold change) was > 0.25 or < −0.25. 14 clusters were visualized with Uniform Manifold Approximation and Projection (UMAP). Each cluster was manually annotated using canonical markers for mammalian mammary gland tissue defined in cattle, mice, and humans [12,28–30]. The visualization of DEGs was performed using ComplexHeatmap (v2.22.0) [31].

### Gene set enrichment analysis

Gene set enrichment analysis (GSEA) was conducted using fgsea package (v1.32.4) [32] based on the hallmark gene sets from MSigDB database [33]. DEGs between HS and TN from each cell cluster were ranked by log2(fold change) from the most upregulated to the most downregulated genes, and the GSEA analysis was conducted with 10,000 permutations to identify the biological pathways affected by HS.

### CellChat analysis

The cell-cell interaction analysis was conducted according to the CellChat (v1.6.1) [34]. Specifically, the CellChat object was created from the Seurat object for each sample, and all CellChat objects were merged for the comparative analysis. The differential cell-cell signaling interaction strength among clusters was calculated and visualized using the netVisual_heatmap function. The ligand-receptor interactions were visualized with the netVisual_bubble function, while the communication probability of ligand-receptor pairs between clusters for each ligand-receptor pair was estimated using the computeCommunProb function.

### Pseudotime trajectory analysis

The pseudotime trajectory analysis was conducted according to the Monocle3 (v1.4.26) [35] following the developer’s tutorial on TN and HS samples. Luminal 1, Luminal 2, and Luminal AV cell clusters were selected for trajectory mapping, with the Luminal 2 cluster designated as the undifferentiated root. Differential gene expression along the pseudotime trajectory was analyzed using the graph_test function to identify trajectory-dependent genes (q value < 0.05). Within each sample, trajectory-dependent genes were grouped into three modules by unsupervised hierarchical clustering and visualized using the ComplexHeatmap package (v2.22.0) [31]. Functional enrichment analyses of genes from each module were performed with the clusterProfiler package (v 4.14.6) [36] to identify overrepresented biological processes and pathways.

### Regulatory network inference and clustering (SCENIC) analysis

Gene regulatory networks were inferred using the SCENIC package (v1.1.2) with associated packages [37]. Transcription factor (TF)-target co-expression modules and gene regulatory networks were first identified using the GENIE3 algorithm, and candidate regulons were then refined with RcisTarget motif enrichment analysis. Single-cell regulon activity scores were computed using the AUCell algorithm. Differential regulon activity between HS and TN was tested with a two-sided Wilcoxon rank-sum test and corrected using the Benjamini-Hochberg FDR method. Regulons with FDR < 0.05 were considered significantly altered between groups. Regulon activity heatmaps were visualized using the pheatmap package [38] in R, and selected TF-target DEGs networks were visualized by ggraph (v2.2.2).

## Results

### Heat stress-induced physiological changes in dairy cows

In this study, nine multiparous lactating Holstein cows (n = 3 per treatment) were housed in respiration chambers and assigned to one of three environmental conditions: thermoneutrality (TN), heat stress (HS), and thermoneutrality with pair-feeding (PF) (**Fig. 1**). One cow in the PF group was removed due to health issues, resulting in a final sample size of eight cows. Environmental conditions were determined by temperature-humidity index (THI) (**Additional file 1: Fig. S1**). Compared with TN and PF cows, HS cows showed increased skin and rectal temperatures as well as elevated heart and respiration rates during each treatment day (**Fig. 2A-D**). Rectal temperatures increased by up to 1.9°C in HS cows compared to TN cows. Peak rectal (40℃) and skin (37.2℃) temperatures were observed in the afternoon and remained at similar levels during the evening measurement (**Additional file 2: Table S1**). There was no difference in the vaginal temperature among the treatments during the baseline period (*P* = 0.32). HS cows had a greater vaginal temperature than both TN and PF cows during the study period (*P* = 0.05) (**Fig. 2E**).

**Fig. 2.**
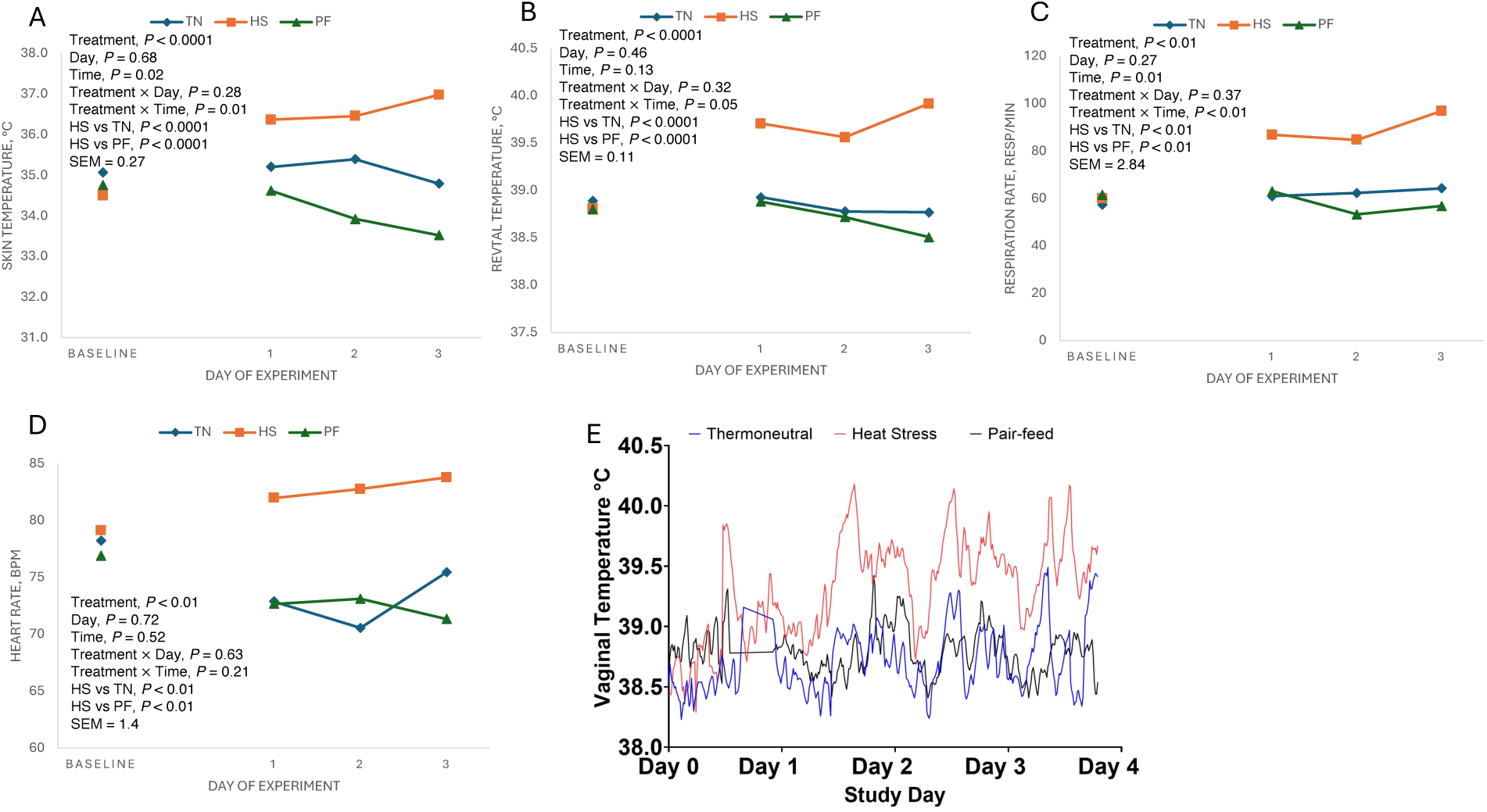
Clinical assessments of dairy cows under environmental stress. Effects of environmental conditions on (A) skin temperature, (B) rectal temperature, (C) respiration rate, (D) heart rate, and (E) vaginal temperature of lactating Holstein cows. BL, baseline values during the acclimation period.

Dry matter intake was lower in HS cows than in TN cows (*P* = 0.04), whereas PF cows had a similar DMI to HS cows, as expected from the experimental design (*P* = 0.93) (**Table 1**). Water intake was greater in HS cows than both TN and PF cows (*P* = 0.01). Heat stress decreased milk yield, 3.5% Fat Corrected Milk (FCM), and Energy Corrected Milk (ECM) compared with TN cows (*P* < 0.01). Efficiencies of milk, 3.5% FCM, and ECM were significantly reduced in HS cows compared with PF cows (P < 0.05) but were similar to those of TN cows. Milk fat, protein, lactose and total solids percentage did not differ among treatments. However, HS cows produced lower yields of milk fat, protein and lactose compared to TN cows (*P* < 0.05), with protein yield also lower than PF cows (*P* = 0.03). The MUN was lower in HS cows compared to PF cows (*P* = 0.03) but higher than TN cows (*P* = 0.01).

**Table 1.** Effects of environmental conditions on productive performance, milk composition, milk solids, and feed efficiency in lactating dairy cows.

| Variable | Treatment<sup>1</sup> |  |  |  | P-value<sup>2</sup> |  |  |  |  |
| --- | --- | --- | --- | --- | --- | --- | --- | --- | --- |
|  | TN | PF | HS | SEM | Treatment | Day | Treatment x Day | HS vs TN | HS vs PF |
| **Productive performance** |  |  |  |  |  |  |  |  |  |
| DMI, kg/d | 18.53 | 13.25 | 12.98 | 1.74 | <0.01 | 0.03 | 0.72 | 0.04 | 0.93 |
| Water intake, L/d | 50.87 | 63.26 | 99.56 | 9.24 | 0.02 | 0.59 | 0.39 | 0.01 | 0.01 |
| Milk Yield, kg/d | 19.39 | 17.13 | 14.33 | 1.37 | <0.01 | 0.02 | 0.41 | <0.01 | 0.07 |
| 3.5% FCM, kg/d | 27.04 | 22.62 | 21.09 | 1.60 | 0.02 | 0.14 | 0.96 | 0.01 | 0.54 |
| ECM, kg/d | 26.48 | 22.47 | 20.65 | 1.47 | 0.01 | 0.11 | 0.92 | <0.01 | 0.43 |
| **Milk composition, %** |  |  |  |  |  |  |  |  |  |
| Fat | 6.48 | 4.50 | 5.54 | 0.63 | 0.15 | 0.66 | 0.39 | 0.09 | 0.11 |
| Protein | 3.92 | 3.70 | 3.75 | 0.40 | 0.89 | 0.68 | 1.00 | 0.77 | 0.94 |
| Lactose | 4.59 | 4.51 | 4.79 | 0.03 | 0.10 | 0.14 | 0.20 | 0.14 | 0.07 |
| TS | 15.62 | 16.06 | 15.75 | 0.49 | 0.76 | 0.62 | 0.57 | 0.74 | 0.56 |
| SCS | 3.69 | 3.41 | 3.13 | 0.36 | 0.68 | 0.60 | 0.91 | 0.48 | 0.69 |
| MUN, mg/dL | 7.85 | 12.91 | 10.57 | 0.65 | <0.01 | 0.23 | 0.87 | 0.01 | 0.03 |
| **Milk solids, kg/d** |  |  |  |  |  |  |  |  |  |
| Fat | 1.14 | 1.01 | 0.90 | 0.10 | 0.07 | 0.15 | 0.97 | 0.03 | 0.43 |
| Protein | 0.73 | 0.71 | 0.56 | 0.04 | <0.01 | 0.02 | 0.30 | <0.01 | 0.03 |
| Lactose | 0.94 | 0.81 | 0.64 | 0.05 | <0.01 | 0.02 | 0.29 | <0.01 | 0.06 |
| **Feed efficiency ratio** |  |  |  |  |  |  |  |  |  |
| Milk<sup>3</sup> | 1.08 | 1.44 | 1.07 | 0.08 | <0.01 | 0.84 | 0.93 | 0.88 | <0.01 |
| 3.5% FCME<sup>4</sup> | 1.53 | 2.00 | 1.52 | 0.16 | 0.07 | 0.88 | 0.99 | 0.94 | 0.05 |
| ECME<sup>5</sup> | 1.51 | 1.98 | 1.48 | 0.15 | 0.05 | 0.87 | 0.99 | 0.88 | 0.03 |

<sup>1</sup>Nine pregnant multiparous and lactating Holstein cows were randomly assigned to one of three environmental treatments: thermoneutrality (TN; n = 3, THI = 68), heat-stress (HS; n = 3, THI = 74- 86), thermoneutrality but pair-fed to HS (PF, n = 3, THI = 68).
<sup>2</sup>Statistical significance was considered when $P \leq 0.05$ . Trend towards significance was considered when $0.05 < P \leq 0.15$ .
<sup>3</sup>Milk efficiency was calculated as the ratio of milk yield to DMI.
<sup>4</sup>ECM efficiency was calculated as the ratio of ECM to DMI.
<sup>5</sup>3.5% FCM efficiency was calculated as the ratio of 3.5%FCM to DMI.

### Identification of cell types in the bovine mammary gland using snRNA-seq

We performed snRNA-seq on samples collected from TN, HS, and PF cows to characterize cell populations in the bovine mammary gland under different environmental conditions. After quality control filtering based on the number of detected genes (nFeature_RNA), total read counts (nCount_RNA), and mitochondrial and ribosomal percentage, 5,201, 7,711, and 1,679 high-quality nuclei were retained from TN, HS, and PF samples, respectively (**Additional file 4: Fig. S2**).

Dimensionality reduction and clustering were conducted using UMAP, which identified 14 transcriptionally distinct clusters (**Fig. 3A**). To assign biological identities to these clusters, we did manual annotation using canonical marker genes from previously published mammary gland single-cell datasets [12,28–30] (**Fig. 3B-D and Additional file 5: Fig. S3**). Based on marker gene expression, six clusters were identified as epithelial, four as stromal, and four as immune cell population (**Fig. 3A-D**). Among all identified clusters, epithelial cells formed the major population on the UMAP, including four luminal and two myoepithelial clusters (**Fig. 3A**). Luminal 1, Luminal 2, and Luminal AV clusters were positioned close to each other on the UMAP, forming the largest epithelial population, while Luminal HS cells were spatially separate from these three clusters. Myoepithelial cells formed the second largest group, whereas immune and stromal cells occupied distinct regions on the UMAP (**Fig. 3A**).

**Fig. 3.**
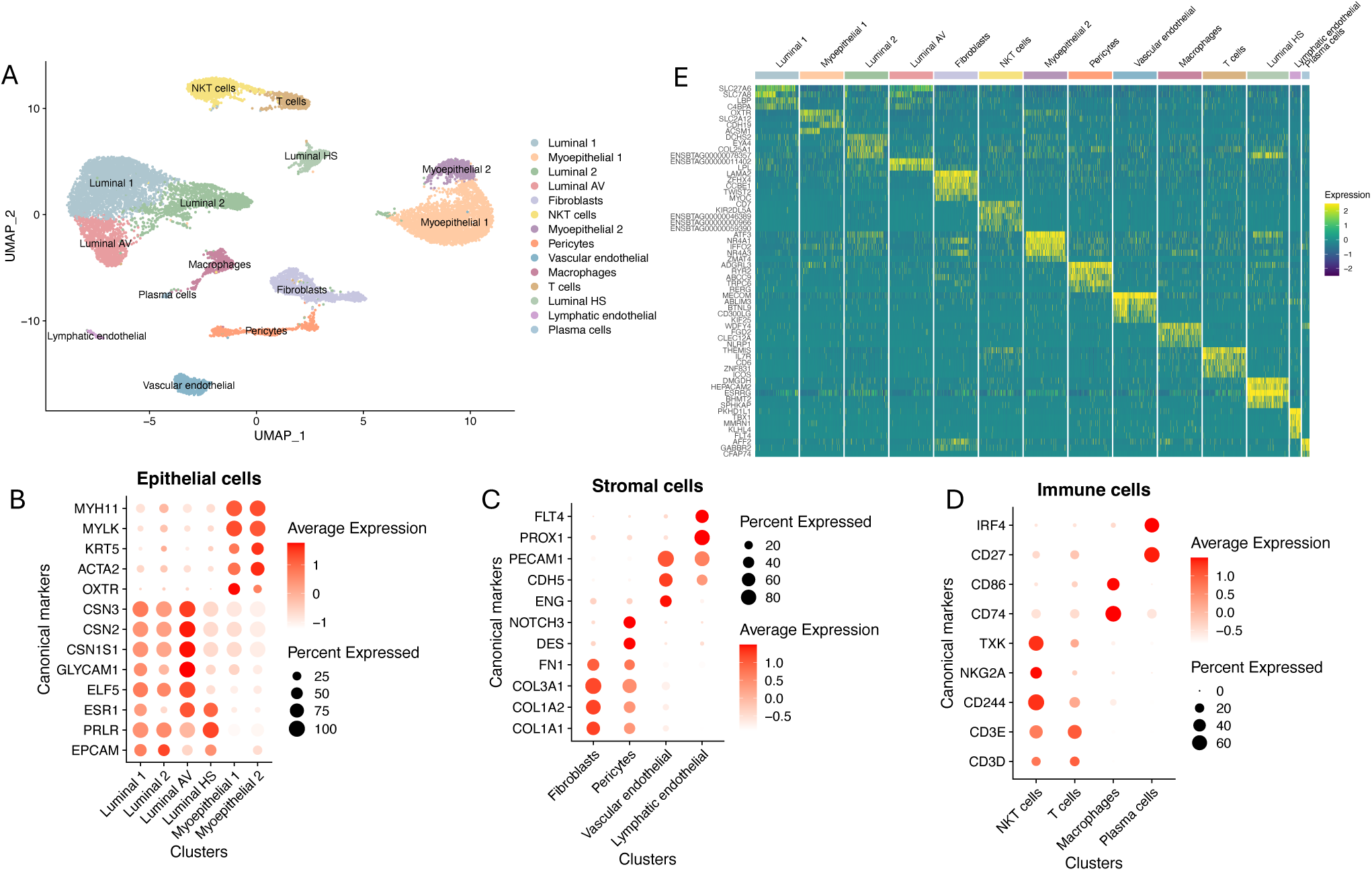
Identification of bovine mammary gland cells by snRNA-seq. **A.** UMAP visualization of all mammary gland nuclei colored by identified cell clusters. **B-D.** Dot plots were used to visualize the expression of canonical markers of each cell cluster within (B) epithelial, (C) stromal, and (D) immune cells. **E.** Heatmap of cell type-specific gene signatures. Luminal AV, luminal alveolar; Luminal HS, luminal hormone-sensing; NKT, natural killer T cells.

Among epithelial clusters, two luminal clusters expressed markers of alveolar (*GLYCAM1*, *CSN3*, and *ELF5*) and hormone-sensing (*ESR1* and *PRLR*) lineages and were designated as Luminal AV and Luminal HS, respectively (**Fig. 3B)**. Two additional luminal clusters co-expressed markers of both lineages and were annotated as Luminal 1 and Luminal 2. Such mixed populations are uncommon but have been previously reported in mouse mammary single-cell dataset [12]. Myoepithelial cells expressed markers such as *KRT5*, *ACTA2*, and *OXTR*.

Among stromal populations, fibroblasts expressed *COL1A1 and FN1*, pericytes expressed *DES* and *NOTCH3*, vascular endothelial expressed *ENG* and *CDH5*, and lymphatic endothelial cells expressed *PROX1* and *FLT4* (**Fig. 3C**). The immune population included natural killer T (NKT) cells expressing *CD244* and *TXK*, macrophages expressing *CD74* and *CD86*, T cells expressing *CD3D* and *CD3E*, and plasma cells expressing *CD27* and *IRF4* (**Fig. 3D)**.

In addition, we identified top expressed genes in each of the 14 cell clusters (**Fig. 3E**). For example, *SLC27A6* and *SLC7A8* were highly expressed in Luminal 1, while *DCHS2* and *COL25A1* were highly expressed in Luminal 2. These genes may serve as novel markers for distinguishing heterogeneous cell types within the mammary cellular population. Collectively, this annotation defined 14 transcriptionally distinct cell clusters within the bovine mammary gland, encompassing major epithelial, stromal, and immune cell populations, thereby establishing a comprehensive single cell type reference for subsequent analyses of heat stress-induced transcriptional changes. The PF sample was excluded from downstream analyses due to its low cell count, particularly in the Luminal HS cluster (cell number = 2) and lymphatic endothelial cell cluster (cell number = 2) (**Additional file 6: Fig. S4 and Additional file 7: Table S3 ).**

### Cell type-specific responses to HS

To examine how different mammary cell populations respond to thermal stress, we compared the transcriptional profiles between TN and HS conditions. Cluster-wise differential expression analysis revealed large variability in transcriptional responsiveness among cell types. The Luminal 1 cluster exhibited the highest number of 3,257 differentially expressed genes (DEGs), and lymphatic endothelial cells showed the lowest number of 3 DEGs (**Fig. 4A** and **Additional file 8: Table S4**). Under HS, several epithelial clusters showed decreased expression of *VNN2*, *SLC35F3*, and *ACTC1*, suggesting reduced cellular integrity, metabolic activity, and contractile function (**Fig. 4A**). In immune cells, *TMC5* and *IL2RA* were downregulated, indicating a potential dampening of immune signaling. Within stromal populations, *MLIP* and *CYP1A1* expression was reduced, consistent with impaired cellular homeostasis and metabolism. In contrast, *NOTCH3*, *TUFT1*, and *COL25A1* showed increased expression in epithelial cells under heat stress, suggesting enhanced structural remodeling under stress. Additionally, *CCN1*, *SPP1*, and *ITGB8* were elevated in immune and stromal cells, reflecting an activated inflammatory response and disrupted cellular development under thermal stress (**Fig. 4A**).

**Fig. 4.**
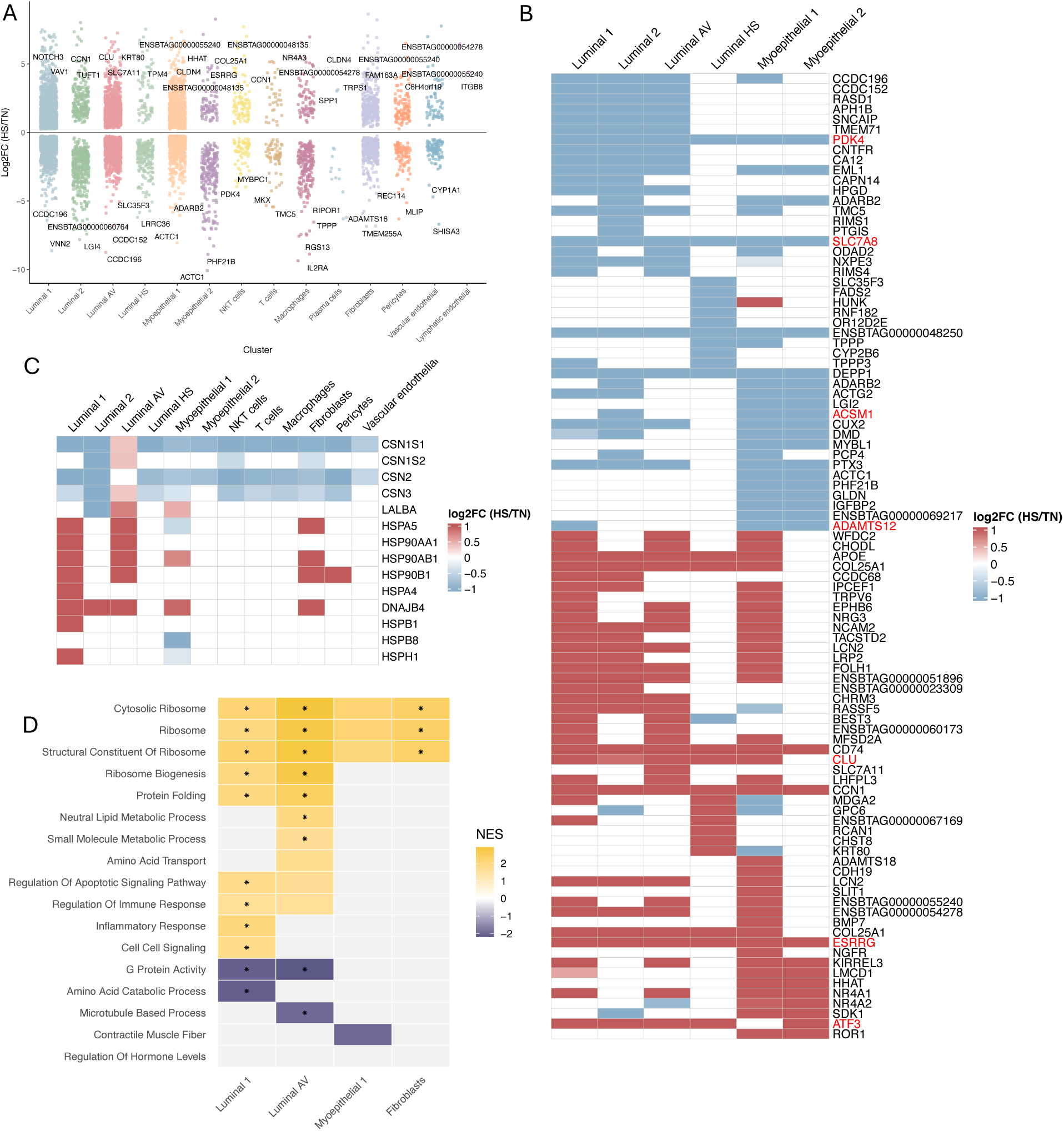
Gene expression profiles of mammary cells under HS. **A.** Volcano plot showing differentially expressed genes (DEGs) identified in each cluster between HS and TN conditions. **B.** Heatmap showing top DEGs under HS in epithelial clusters. **C.** Heatmap showing expression of milk protein genes and heat shock protein genes in each cluster. **D.** Hallmark terms enriched under TN or HS conditions in HS-responsive clusters obtained by GSEA.

Top HS-induced DEGs in epithelial cells were shown in **Fig. 4B**. Genes associated with metabolic regulation and tissue remodeling were significantly reduced in expression upon HS, including *PDK4*, *SLC7A8*, *ACSM1*, and *ADAMTS12,* reflecting the impact of HS on these critical biological processes. In contrast, HS triggered a profound transcription shift toward stress-adaptive and cellular survival pathways, with increased expression of *CLU*, *ESRRG, SLC7A11,* and *ATF3.* These genes are known regulators of protein folding, energy metabolism, oxidative response, and cellular stress response, indicating that HS imposes substantial proteotoxic and metabolic stress on mammary epithelial cells.

Compared to TN conditions, milk protein genes (*CSN1S1*, *CSN2*, and *CSN3*) showed significantly reduced expression under HS, except for a mild increase observed in Luminal AV cells (**Fig. 4C**), indicating cell type-specific HS responses in milk protein gene regulation. In contrast, heat shock protein (HSP) genes (*HSPA5*, *HSP90AA1*, *HSP90AB1, HSP90B1,* and *DNAJB4*) were significantly upregulated under heat stress, particularly in Luminal 1, Luminal AV, Myoepithelial 1, and Fibroblast clusters (**Fig. 4C**). Interestingly, Luminal 2 and Luminal HS cells exhibited limited or no elevation in HSP gene expression. Similarly, other stromal and immune cell populations showed minimal HS-induced HSP activation, except for an increased *HSP90B1* expression observed in pericytes.

Gene set enrichment analysis (GSEA) on HS-responsive clusters revealed significant enrichment of multiple ribosome-related pathways (**Fig. 4D**), including cytosolic ribosome and ribosome biogenesis, indicating increased translational demand and cellular attempts to restore protein homeostasis. In Luminal 1, HS induced enrichment of apoptotic signaling and inflammatory response, consistent with a stress-activated cellular state. Importantly, the major milk-synthesizing population, Luminal AV cells, showed significant enrichment of protein folding and metabolic processes, suggesting efforts to preserve secretory capacity under challenge. In contrast, G protein activity was suppressed under HS, indicating disruption in lactation and hormone regulation (**Fig. 4D**). This pattern suggests that heat stress compromises the normal functions of luminal cells and redirects cellular resources toward maintaining proteostasis rather than milk protein synthesis. Collectively, these findings reveal that HS disrupts the transcriptional and metabolic equilibrium of the mammary epithelium, impairing its capacity to sustain efficient milk production under thermal stress.

### Mammary gland cell-cell interaction under TN and HS

Cell-cell communication analysis revealed a clear HS-dependent remodeling of cellular signaling. Signaling pathways enriched under TN and HS conditions were ranked by their overall strength (**Additional file 9: Fig. S5A**). Pathways related to mammary development and differentiation such as *WNT, NEGR*, and *BMP* showed strong communication strength under TN, whereas pathways such as *SPP1, THBS*, and *IGF* were more active under HS . The change in interaction strength between clusters showed that signaling sending from Fibroblasts to Luminal cells decreased in strength, while Fibroblasts to myoepithelial cells increased under HS compared to TN (**Fig. 5A**). Inter-cluster signaling strength of selected pathways was also shown (**Additional file 9: Fig. S5B-D**). *NRG* and *CADM* pathways were more active under TN and predominantly originated from epithelial cells. In contrast, *SPP1* and *THBS* pathways were more active under HS, with signaling sent from Luminal 1 and Myoepithelial 2 to other epithelial cells, respectively.

**Fig. 5.**
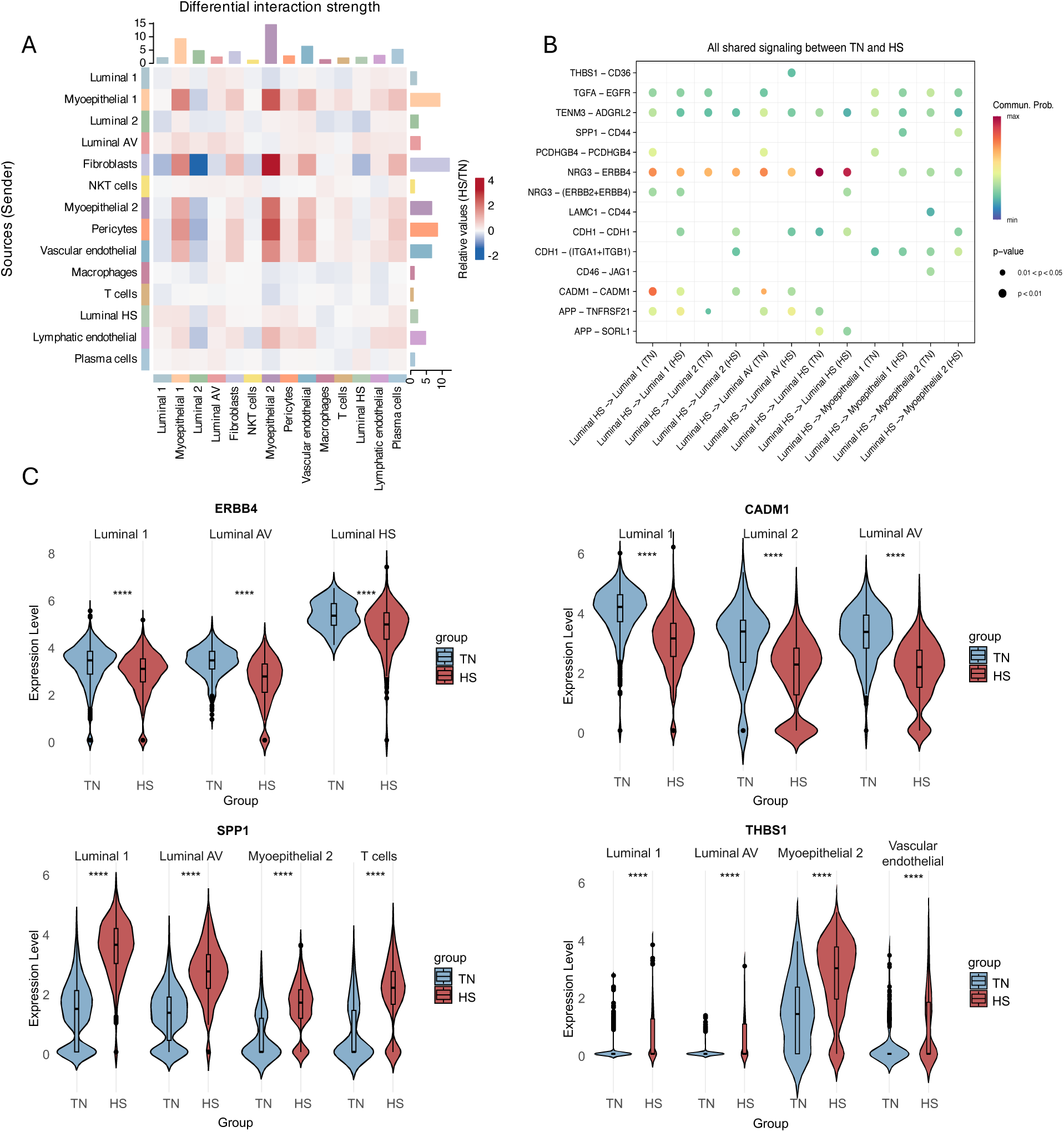
Cell-cell communication under environmental conditions. **A.** Heatmap showing differential interaction strength between clusters under HS. **B.** Dot plot showing ligand-receptor pair signaling under HS and TN conditions. **C.** Violin plots showing differential expression of selected genes under HS and TN conditions.

Given the role of luminal HS cells in receiving hormone cues and transmitting intercellular signals, we further investigated the specific ligand-receptor signals originating from the Luminal HS cluster to other epithelial clusters. We found that NRG3-ERBB4 signaling, associated with milk synthesis, and CADM1-CADM1 signaling, associated with cell adhesion, were enriched under TN. *THBS-CD36* signaling, related to endoplasmic reticulum stress, and *SPP1-CD44* signaling, related to cell proliferation and inflammation, were enriched under HS (**Fig. 5B**). Gene-expression changes were consistent with these patterns, as *ERBB4* and *CADM1* were upregulated under TN in Luminal cells, while *APP* and *THBS1* were upregulated under HS across various clusters (**Fig. 5C**).

### Pseudotime analysis revealed the developmental trajectory of luminal cells

Pseudotime analysis was performed on Luminal clusters, including Luminal 1, Luminal 2, and Luminal AV clusters to reveal the developmental trajectory, with Luminal 2 set as the root due to higher expression of *KIT*, a luminal progenitor marker (**Additional file 10: Fig. S6A**). Luminal HS cluster was excluded because it formed a separate cluster (**Fig. 3A**). Under the TN condition, Luminal AV was mapped as the terminal point of the trajectory (**Fig. 6A and Additional file 10: Fig. S6B**), consistent with its differentiated secretory cell state. Variable genes along pseudotime were analyzed and the top 100 genes were grouped into three modules corresponding to early, mid, and late stages of the trajectory (**Fig. 6B**). Early-stage genes included *LTF* and *TXNIP*, mid-stage genes included *SAT1* and *FOXP1*, while late-stage genes included multiple casein genes. GO analysis of each module showed distinct cellular functions enriched at each stage (**Fig. 6C**). Early stage (module 1) genes were enriched in RNA-related metabolism and regulation, middle stage (module 2) genes were enriched in muscle development, and late stage (module 3) genes were enriched in cellular response to hormone stimulus, consistent with the differentiation trajectory of Luminal cells.

**Fig. 6.**
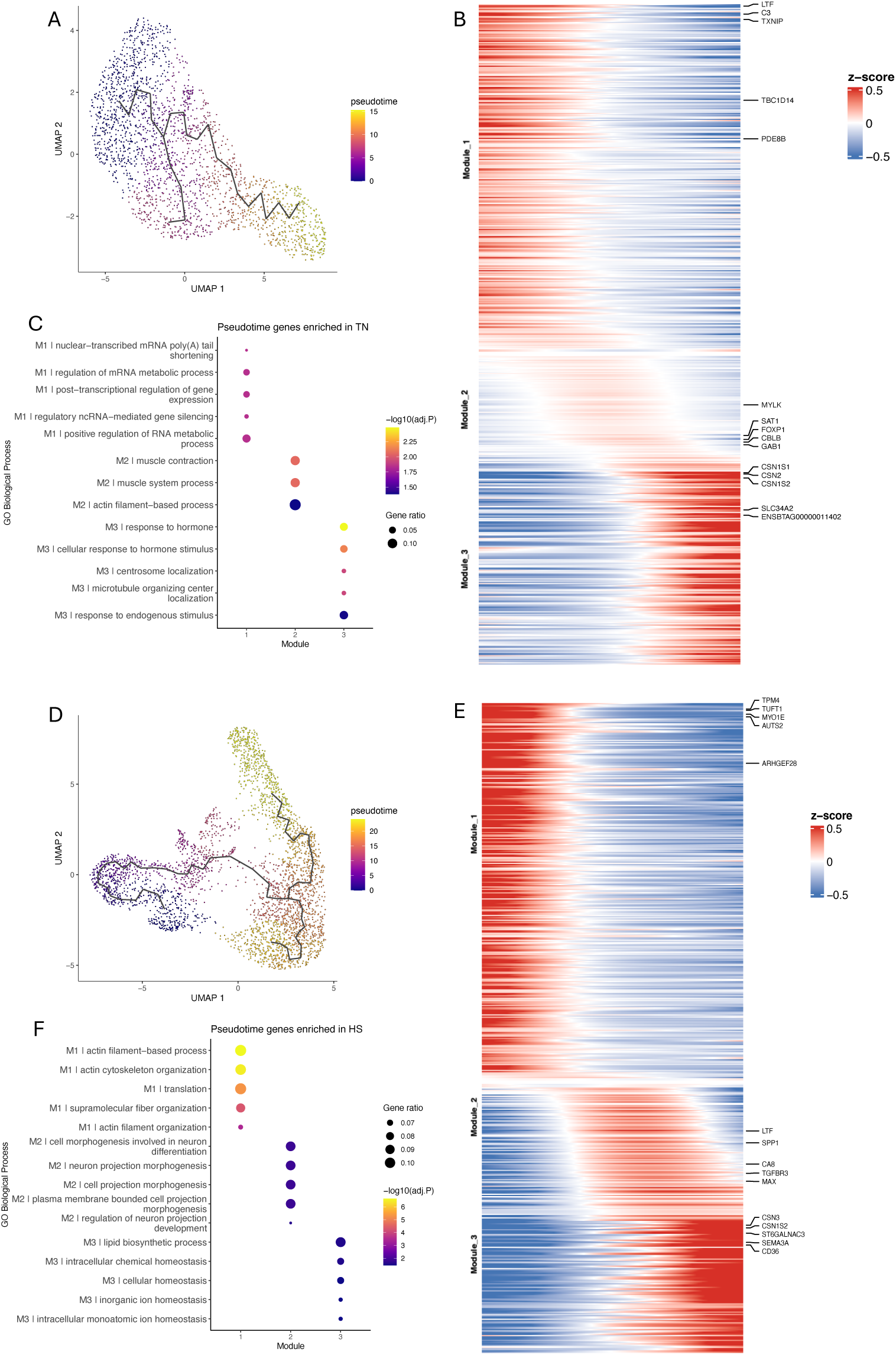
Pseudotime analysis revealed luminal cell developmental trajectory. **A.** pseudotime trajectory of luminal cells under TN. **B.** Heatmap showing genes enriched along pseudotime modules under TN. **C.** GO terms enriched on genes expressed in each module. **D.** Pseudotime trajectory of luminal cells under HS. **E.** Heatmap showing genes enriched along pseudotime modules under HS. **F.** GO terms enriched on genes expressed in each module.

We conducted the same analysis in the HS condition. Luminal AV cells were still mapped as the late stage along pseudotime with the expression of milk protein genes. The enriched functional processes for the late stage (module 3) genes were redirected toward homeostasis processes (**Fig. 6D-F** and **Additional file 10: Fig. S6C**). In contrast, early- and mid-stage gene-expression profiles and GO enrichments closely matched those under TN, indicating that HS-driven remodeling is concentrated in terminal luminal states.

### Transcriptional regulatory networks differ by cell type and are impacted by HS

To profile transcription factor (TF)-target gene regulatory networks in the mammary gland, we quantified regulon activities under TN and HS conditions. Regulons clustered according to cell populations, suggesting cell type-specific transcriptional regulation (**Fig. 7A**). Comparing to TN, EGR1, ATF3, and JUN showed elevated activity under HS, especially in Luminal epithelial cells. In contrast, KLF6, FOXP1, and FOXP2 were repressed by HS. Within myoepithelial cells, PURG and MYBL1 were highly repressed by HS. We next mapped the downstream targets of ATF3 and FOXP2, linking several DEGs regulated by these TFs. Consistently, most targets regulated by HS-activated ATF3, including *RCAN1, SCN5A*, and *DUSP5*, were upregulated under the HS condition, whereas most targets regulated by HS-repressed FOXP2, including *PDK4, RASD1*, and *SNCAIP*, were downregulated in HS condition (**Fig. 7B**).

**Fig. 7.**
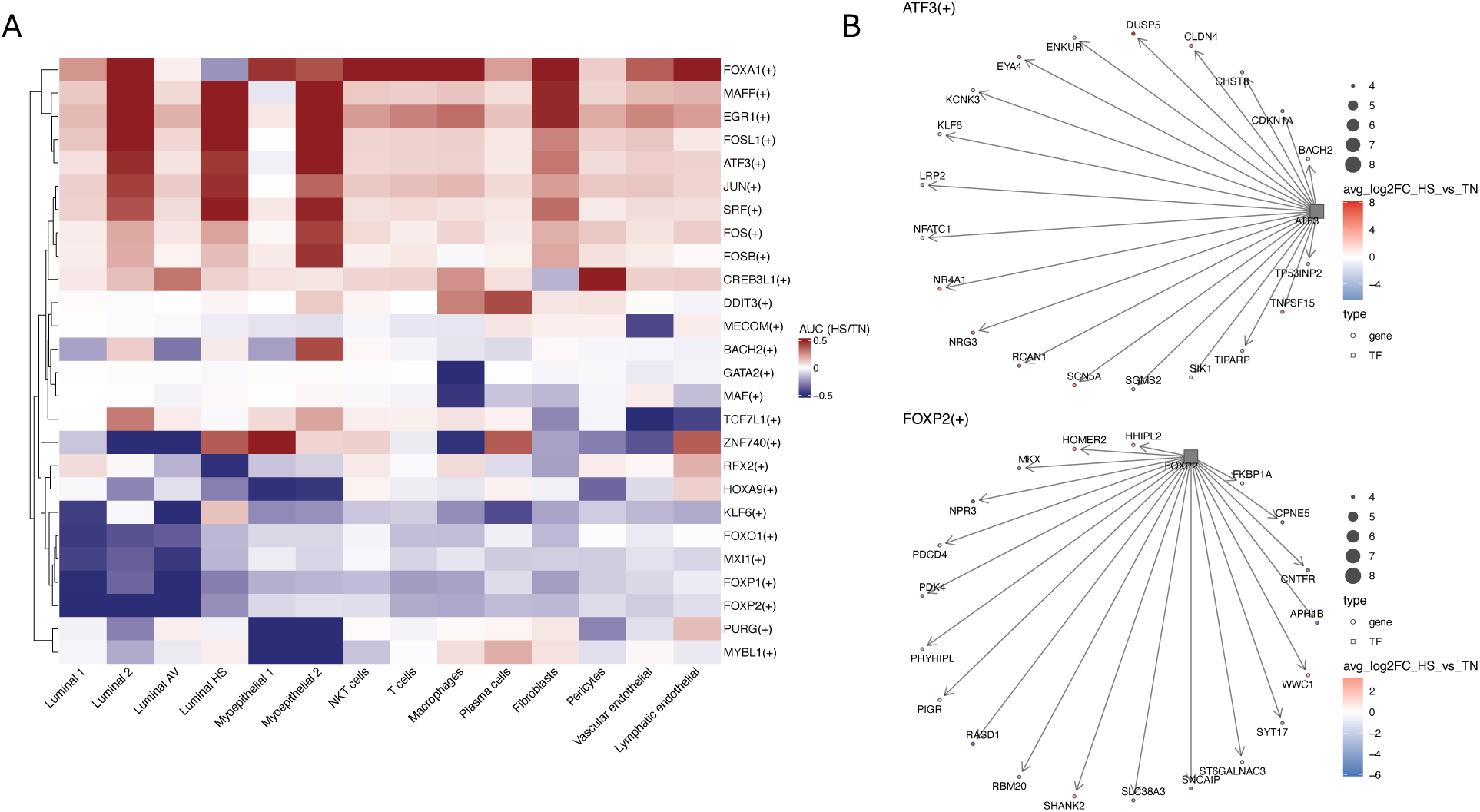
Transcription factor regulatory network inference in mammary cells. **A.** cell type-specific transcription factors (TFs) that were activated under TN or HS. **B.** Network visualization of selected TFs and their downstream target DEGs.

## Discussion

In this study, we combined physiological measurements with single-nucleus RNA sequencing (snRNA-seq) to investigate how heat stress alters dairy cattle production and cellular function of the mammary gland. Although biopsies within each condition were pooled to maximize nuclei recovery, thereby limiting the ability to capture individual variation, our snRNA-seq dataset provides the first atlas of mammary cellular responses to thermal stress. The removal of one PF cow further reduced nuclei input, leading to insufficient cell numbers for downstream analysis. Thus, the PF group was excluded from transcriptomic comparisons. Future work with biological replicates for sequencing will be needed to validate these observations and more clearly separate the effects of metabolic restriction from heat stress.

The thermal environment experienced by dairy cows is commonly evaluated using THI, which reflects the combined effects of relative humidity and ambient temperature [20]. For lactating cows, a THI below 68 is generally regarded as thermoneutral, whereas values exceeding this threshold, especially above 72, indicate a heat stress condition due to cows’ high metabolic heat production [4]. In this study, respiration chambers were used to precisely regulate environmental conditions, with TN and PF cows maintained at a THI of 68 and HS cows exposed to a maximum THI of 86 during the day and 74 at night. Clinical signals of heat stress, including increased skin, rectal, and vaginal temperatures and elevated heart and respiration rates, were observed in HS cows and were consistent with responses reported in our previous heat stress studies [39,40].

Reduced DMI is a characteristic response of dairy cows under heat stress, as an adaptive response to limit metabolic heat production. Earlier work has reported more than 30% DMI reduction [39,41]. In this study, HS cows consumed approximately 29.95% less DMI compared to TN cows. Incorporation of the PF group enabled separation of intake-driven versus heat-stress-driven effects on lactational performance. Comparison among groups indicated that the decreased DMI explained ∼45% of the decline in milk yield, while direct thermal stress effects accounted for the remaining ∼55%. This observation is consistent with previous studies that utilized pair-feeding controls [7,41,42] and underscores the direct impacts of thermal stress on lactation. Increased water intake in HS cows in this study is also consistent with previous reports describing greater water consumption under heat stress to aid heat dissipation and maintain hydration [43,44].

Impairments in milk protein and fat synthesis have been documented in multiple studies. Although fat, protein, and lactose concentrations were not altered, HS cows exhibited reduced yields of all three solids relative to TN cows. The decreased fat yield has been attributed to reduced *de novo* fatty acid synthesis under heat stress [45,46]. As reported previously, the reduction in milk protein yield aligns with downregulation of milk protein genes, particularly caseins, under stress [8,9]. Although a previous study reported a tendency for low lactose yield in HS condition [6], a weak association between HS and lactose yield has been suggested [47]. Therefore, the lower lactose yield in HS cows than TN cows observed in this study might be attributed to reduced DMI and thus lower availability of glucose precursors [48].

MUN is influenced by both dietary crude protein intake and ambient temperature [49]. In this study, HS cows had a higher MUN than TN cows, suggesting a diminished utilization of dietary protein under stress, which has also been reported previously [39]. We also observed a lower MUN in HS cows than in PF cows. Although this finding differs from previous studies [39,41], PF and HS cows consumed comparable feed yet differed in MUN and milk protein yield, indicating that reduced intake alone cannot account for the nitrogen response observed under heat stress. These differences may reflect variation in animal characteristics, such as stage of lactation and production level, as well as variation in heat stress duration and intensity among studies, as a positive relationship between ambient temperature and humidity and MUN concentrations has been proposed [49]. The feed efficiency is a key indicator for evaluating nutrient partitioning under thermal challenge. HS cows showed reduced efficiencies for milk yield, 3.5% FCM, and ECM compared to PF cows. This result suggests a stress-induced decrease in efficiency beyond the effects of low DMI, consistent with previous findings [39,50].

snRNA-seq analysis identified 14 cell clusters, including Epithelial, Immune, and Stromal populations. Within epithelial cells, Luminal alveolar (luminal AV) cells expressed secretory markers such as *GLYCAM1* and *CSN3*, confirming their role in milk synthesis. Luminal hormone-sensing (Luminal HS) cells expressed *ESR1* and *PRLR*, consistent with their function as hormone receptors [29]. Luminal 1 and Luminal 2 clusters expressed markers of both lineages, suggesting a transitional or an undifferentiated state [12]. Luminal 1 highly expressed *SLC27A6* and *SLC7A8*, which are associated with lipid and amino acid metabolism in the mammary gland [51,52], whereas Luminal 2 expressed *DCHS2*, which is related to cell adhesion [53].

HS profoundly altered the molecular functions of mammary epithelium. Consistent with the observed reduction in milk protein yields, *CSN1S1*, *CSN2*, and *CSN3* were broadly downregulated in epithelial cells under HS. In contrast, heat shock protein (HSP) genes were strongly induced in epithelial cells and fibroblasts. HSP genes act as molecular chaperones to maintain proper protein folding under stress, and their induction in milk-synthesizing cells suggests a cellular response to proteotoxic stress that may directly compromise milk secretion [54]. Among HS-responsive epithelial clusters, the expression of *CLU*, *ESRRG,* and *SLC7A11* was induced. As a HS chaperone, *CLU* binds to misfolded proteins and prevents apoptosis [55]. *ESRRG* regulates mitochondrial function and energy metabolism, and its upregulation suggests increased energy need under HS [56]. *SLC7A11* participates in the synthesis of glutathione, an important antioxidant against oxidative stress [57].

GSEA results further revealed a strong enrichment of pathways related to the ribosome in HS-responsive cells. Ribosomes translate mRNA into proteins, and their activation may reflect an elevated proteostasis demand to maintain basic cellular functions [58]. Protein folding and metabolic processes were enriched in Luminal AV cells, and, interestingly, Luminal AV cells upregulated both milk protein genes and HSP genes, suggesting that these cells may experience both metabolic and proteostatic stress as they attempt to maintain secretory activity under thermal stress [59,60]. Moreover, fibroblasts, which mediate extracellular matrix (ECM) remodeling, upregulated *DUSP5* under HS, a regulator of stress recovery pathways [61]. Together, these findings highlight a cell type-specific response to thermal stress and an overall increased proteostasis demand across cells.

Cell-cell communication between stromal and epithelial cells was markedly altered by heat stress. Signaling from Fibroblasts to Luminal cells was reduced under HS, whereas their communication to myoepithelial cells was enhanced. Fibroblasts serve as a major source of paracrine cues that regulate epithelium, and, although their roles under HS have not been well characterized, fibroblast-derived signals are known to influence inflammation, epithelial-mesenchymal transition (EMT), or epithelial differentiation under various conditions [62,63]. Therefore, the HS-induced changes in fibroblast-epithelial signaling observed in our study may be a potential molecular mechanism through which stromal signals impair milk synthesis.

Within epithelial cells, Luminal HS cells receive hormonal cues and transmit signals to other cells to regulate mammary function. To further examine how thermal stress influences epithelial communication, we profiled ligand-receptor pairs with altered signaling strength. *ERBB4*, which binds to STAT5 and activates milk synthesis pathways [64], and *CADM1*, which mediates cell adhesion and maintains epithelium structure [65], were both downregulated under HS. In contrast, *SPP1* is known as a positive regulator of milk protein gene expression [66], and its upregulation under HS in luminal cells, particularly Luminal AV cells, may represent a compensatory mechanism to preserve milk synthesis. Beyond its role in lactation, *SPP1* has been reported to disrupt T cell function under disease or stress conditions [67]. In addition, *THBS1* was also induced by HS and has been implicated as a mediator for endoplasmic reticulum (ER) stress [68]. Overall, these findings reveal that HS reshapes mammary cellular signaling networks, particularly inter-and intra-epithelial communication.

The trajectory analysis revealed a progenitor-to-secretory differentiation of Luminal cells under the TN condition. However, HS redirected the developmental trajectory toward a homeostatic state. Similar disruptions in luminal cell and mammary gland function under HS have been reported in both *in vivo* and *in vitro* studies [59,69,70]. Under TN condition, *LTF* was expressed at the early stage along the pseudotime, whereas under HS its expression shifted to the mid stage. *LTF* is known as a marker for secretory alveolar cells, and its delayed expression under HS may imply a postponed activation of milk synthesis [12]. At the late stage under HS condition, in addition to casein genes, *SEMA3A* and *CD36* were also expressed. *SEMA3A* has been implicated in anti-inflammation and antioxidant responses [71], and *CD36* participates in lipid metabolism and contributes to homeostasis [72]. Functional enrichment analysis of variable genes along pseudotime further revealed the transition to homeostasis at the late stage under HS. These data suggest that HS fundamentally alters Luminal cell fate by delaying secretory differentiation and promoting stress adaptation. Future studies incorporating recovery periods will be important to determine the duration of these developmental trajectory changes.

The transcriptional regulatory network in the mammary gland was also reshaped by thermal challenge. ATF3 is a well-defined stress-induced transcription factor (TF) that is related to glucose metabolism and immune regulation [73], and it was strongly activated under HS, particularly in Luminal 2, Luminal AV, and Myoepithelial 2 clusters. This pattern suggests a strong response to thermal stress in mammary epithelium. Consistently, *NR4A1*, a downstream target of ATF3, was also upregulated under HS and has been previously found in molecular response to heat in dairy cattle [74]. In contrast, FOXP1 and FOXP2 were activated under the TN condition, especially in luminal cells. These TFs are known to regulate mammary ductal morphogenesis and suppress EMT, therefore maintaining epithelial differentiation and structural integrity [75,76]. Target genes of FOXP2, such as *PDK4*, were highly expressed under TN, suggesting an elevated energy demand required for lactation [77]. These altered activities of TFs suggest that HS greatly impacted the regulatory network related to epithelial differentiation and stress response, thus interfering with mammary gland function.

## Conclusions

In this study, we examined the physiological response in dairy cattle and provided the first single-nucleus-resolved transcriptional map of the bovine mammary gland under heat stress. We observed that heat stress significantly reduced DMI, milk yield, milk component yields (fat, protein, and lactose), while water and MUN were increased by heat stress. Our snRNA-seq data show that heat stress markedly altered epithelial cellular functions, cell-cell communication, developmental trajectory, and key transcriptional regulators, potentially contributing to the overall decline in milk production under HS. Overall, these results revealed insights into the cell type-resolved molecular mechanisms underlying heat-induced lactation decline and the strategies to mitigate thermal stress in the dairy industry.

## List of abbreviations

TN: Thermoneutrality HS: Heat stress
PF: Pair feeding
GLYCAM1: Glycosylation-dependent cell adhesion molecule 1
CSN3: Kappa-casein
SLC27A6: Solute carrier family 27 member 6
SLC7A8: Solute carrier family 7 member 8
ESR1: Estrogen receptor 1
PRLR: Prolactin receptor
DUSP5: Dual specificity phosphatase 5
DCHS2: Dachsous cadherin-related 2
CLU: Clusterin
ESRRG: Estrogen-related receptor gamma
SLC7A11: Solute carrier family 7 member 11
ERBB4: Erb-b2 receptor tyrosine kinase 4
CADM1: Cell adhesion molecule 1
SPP1: Secreted phosphoprotein 1
THBS1: Thrombospondin 1
PDK4: Pyruvate Dehydrogenase Kinase 4
FOXP1: Forkhead box P1
FOXP2: Forkhead box P2
NR4A1: Nuclear receptor subfamily 4 group A member 1
ATF3: Activating transcription factor 3
SEMA3A: Semaphorin 3A
CD36: Platelet glycoprotein 4
LTF: Lactotransferrin

## Declarations

### Ethics approval and consent to participate

All experimental procedures were approved by the Cornell University Institutional Animal Care and Use Committee (#2022-0186).

### Consent for publication

Not applicable.

### Availability of data and material

Single-nucleus RNA-seq data have been deposited at NCBI GEO database (GSE317482). This paper does not produce original code. Any additional information required to reanalyze the data reported in this paper is available from the lead contact upon request.

### Competing interests

The authors declare that they have no competing interests.

## Funding

This research was supported by a startup grant from the College of Agriculture and Life Sciences to the Duan lab research. This work was funded by the USDA Federal Capacity Hatch Fund from Cornell Agricultural Station to J.E.D. and J.M.

## Authors’ contributions

X.Y. and S. performed the animal HS experiment. S. enrolled animals, implemented the feeding regimen, led the undergraduate team in daily animal care at LARTU, and collected and analyzed physiological data, while S.P. assisted with data collection, including intake, and milk production. D.C.C., M.M.F., and A.Z. performed mammary gland biopsy and collected liver and blood samples under F.A.L.Y.’s supervision. X.Y. assisted with animal physiological data collection and mammary gland biopsy. Y. F., G.L. shared single-cell data analysis pipelines. X.Y performed the 10x RNA sample process and single nucleus RNA-seq analysis, drafted the manuscript, and designed the figures under JED’s supervision. N.S. managed the animal HS experiment and coordinated with Cornell University Ruminant Center on animal usage. F.L., JWM, and JED designed the project and worked with IACUC. All authors read, revised, and approved the final manuscript.

## Acknowledgements

The authors acknowledge members and undergraduates from Precision Livestock Health (Leal Yepes lab), McFadden lab, and Duan lab for assistance with sample collection. The authors acknowledge all staff at LARTU and Greg Johnson, director of operations at CURC, for help with animal feed.

## Additional files

**Additional file 1: Fig. S1. Environmental conditions in respiration chambers during the experimental period. A-C.** Data were averaged hourly throughout the experiment for (A) ambient temperature, (B) relative humidity, and (C) temperature-humidity index (THI). BL, baseline values during acclimation period; HS, heat stress; TN, thermoneutrality; PF, thermoneutrality and pair-fed.

**Additional file 2: Table S1. Effects of heat stress on clinical assessments of lactating multiparous Holstein cows.** Rectal temperature, skin temperature, respiration rate, and heart rate were measured three times daily (0700, 1200, and 1900 h) in cows maintained under TN, HS, or PF conditions.

**Additional file 3: Table S2. Summary of sequencing and quality control metrics for snRNA-seq libraries.** Summary statistics of sequencing depth and quality control metrics for nuclei from TN, HS, and PF samples.

**Additional file 4: Fig. S2. Quality control metrics for snRNA-seq data. A-C.** Quality control of snRNA-seq data for (A) TN, (B) HS, and (C) PF samples.

**Additional file 5: Fig. S3. Feature plots of marker gene expression.** Feature plots showing expression of marker genes for identified clusters.

**Additional file 6: Fig. S4. Cell proportion and counts across samples. A.** Proportion of each cell cluster in individual samples. **B.** Total cell counts in TN, HS, and PF samples.

**Additional file 7: Table S3. Cell counts per cluster and condition.** Number of cells in each cluster in TN, HS, and PF conditions.

**Additional file 8: Table S4. Differentially expressed genes identified in the HS vs. TN comparison.** List of differentially expressed genes identified between HS and TN conditions.

**Additional file 9: Fig. S5. Signaling pathways enriched in TN or HS conditions**. **A.** Signaling pathways enriched in TN or HS conditions. **B-E.** Intercellular communication networks for selected signaling pathways among identified clusters. NRG, neuregulin; CADM, cell adhesion molecules; SPP1, secreted phosphoprotein 1; THBS, thrombospondin.

**Additional file 10: Fig. S6. Trajectory analysis of luminal clusters. A.** Dot plot showing expression of luminal progenitor marker genes in Luminal 1, Luminal 2, and Luminal AV clusters. **B-C.** Trajectory maps of luminal cells with clusters labeled in (B) TN condition and (C) HS condition. KIT, KIT proto-oncogene, receptor tyrosine kinase; ALDH1A3, aldehyde dehydrogenase 1 family member A3; CD14, monocyte differentiation antigen CD14.

## Notes

### Competing Interest Statement

The authors have declared no competing interest.

